# Monitoring Anti-*Pythium insidiosum* IgG Antibodies and (1->3)-β-D-Glucan in Vascular Pythiosis

**DOI:** 10.1101/300764

**Authors:** Navaporn Worasilchai, Nitipong Permpalung, Pakawat Chongsathidkiet, Asada Leelahavanichkul, Alberto Leonel Mendoza, Tanapat Palaga, Rangsima Reantragoon, Malcolm Finkelman, Pranee Sutcharitchan, Ariya Chindamporn

## Abstract

**Background:** Despite aggressive treatment, vascular pythiosis has a mortality rate of 40%. This is due to the delay in diagnosis and lack of effective monitoring tools. To overcome this drawback, serum beta-D-glucan (BG) and *P*. *insidiosum* specific antibody (*Pi*-Ab) were examined as potential monitoring markers in vascular pythiosis.

**Methods:** A prospective cohort study of vascular pythiosis patients was carried on during January 2010-July 2016. Clinical information and blood samples were collected and evaluated by the BG and *Pi*-Ab assays. Linear mixed effect models were used to compare BG and *Pi*-Ab levels. The *in vitro* susceptibility test was performed in all *P*. *insidiosum* isolates from culture positive cases.

**Results:** A total of 50 patients were enrolled; 45 survived and 5 died during follow-up. The survivors had a significantly shorter time to medical care (p<0.0001) and a significantly shorter waiting time to the first surgery (p<0.0001). There were no differences in BG levels among the groups at diagnosis (p=0.17); however, BG levels among survivors were significantly lower than the deceased group at 0.5 months (p<0.0001) and became undetectable after 3 months. Survivors were able to maintain ELISA value (EV) of *Pi*-Ab above 8, whereas EV among deceased patients was less than 4. *In vitro* susceptibility results revealed no synergistic effects between itraconazole and terbinafine.

**Conclusions:** This study showed that BG and *Pi*-Ab were potentially valuable markers to monitor the disease after treatment initiation. An unchanged BG level at 2 weeks after surgery should prompt an evaluation for residual disease.

## Introduction

Vascular pythiosis is the most common manifestation of human pythiosis caused by *Pythium insidiosum (P*. *insidiosum)*. Vascular pythiosis almost always occurs in patients with underlying hemoglobinopathy complicated by hemochromatosis (1-4). Despite state-of-the-art treatments, the mortality rate of vascular pythiosis is still at 40% (1) and its morbidity, among survivors, is severe due to the need for aggressive amputation to remove all infected tissues (2).

Currently, the following diagnostic criteria are used (1, 2). Presence of typical pathological features, successful isolation of *P*. *insidiosum* (2), positivity for *P*. *insidiosum* specific antibody (*Pi*-Ab) by western blot (5) or enzyme-linked immunosorbent assay (ELISA) (6), and positive PCR using internal transcribed spacer and cytochrome oxidase II (PCR-ITS/*COX2*) primers (7-9). Positive culture in conjunction with zoospore production remains the gold standard (10).

Treatment practices vary across different institutions (2, 6). At King Chulalongkorn Memorial Hospital (KCMH), aggressive surgery, systemic antifungal therapy with a combination of itraconazole and terbinafine (ITC/TRB) as well as the adjunctive use of *P*. *insidiosum* immunotherapy (PIAI) have been recommended in all vascular pythiosis patients under our institutional research protocols (2). A year course of ITC/TRB has been recommended based on a case report of *P*. *insidiosum* periorbital cellulitis in a child who was completely cured by ITC/TRB (11). Our previous study, however, revealed no synergistic effect of ITC/TRB for *P*. *insidiosum* isolates in Thailand (2). In clinical practice, susceptibility testing for this pathogen was not routinely performed and antifungal treatment was not affected by the MICs, given no standardized interpretation.

During the one-year treatment course, patient’s history and physical examination at each clinic visit are the main follow-up tools. Culture or detection of *P*. *insidiosum* DNA at each clinic visit is very unlikely to be successful without invasive surgery to obtain infected tissues. Without standard serologic or inflammatory markers, the sensitivity for the detection of early signs of treatment failure or residual disease is lower. This study was conducted as a preliminary characterization to examine the potential of serum β-D-glucan (BG) and *P*. *insidiosum* specific antibody (*Pi*-Ab) as monitoring markers in vascular pythiosis. *In vitro* susceptibility testing against amphotericin B, voriconazole, itraconazole, fluconazole, anidulafungin, caspofungin, and terbinafine was also performed in all *P*. *insidiosum* isolates from culture positive cases.

## Methods

### Study Design

We performed a prospective cohort study of vascular pythiosis patients who received a combination therapy of surgery, systemic antifungal agents, and immunotherapy with PIAI according to our research treatment protocol at KCMH during January 2010-July 2016.

This study was approved by the Chulalongkorn University Institutional Review Board based on the International Guidelines for Human Research Protection as Declaration of Helsinki, the Belmont Report, Council for International Organizations of Medical Sciences (CIOMS) Guideline and International Conference on Harmonization in Good Clinical Practice (ICH-GCP).

### Patients

All proven vascular pythiosis patients over the age of 18 who received treatment with surgery, systemic itraconazole (Sporal^®^), 100 mg three-times daily, systemic terbinafine (Lamisil^®^), 250 mg twice daily, and immunotherapy with PIAI for 1 year course according to the KCMH protocols were enrolled into this study. The definitive diagnosis of vascular pythiosis was confirmed by two of the following accepted diagnostic criteria: 1) successful isolation of *P*. *insidiosum* 2) positive *Pi*-Ab by *in*-*house* ELISA established by KCMH (KCMH-ELISA) 3) positive PCR-ITS/*COX2* either from the isolates or directly from the clinical specimens (7-9). Informed consent was obtained from all patients and the detailed information regarding the current treatment protocol including PIAI was provided as an investigational treatment under the research protocol.

### Serum and Clinical Data Collection

Eligible patients were enrolled to the study at the time of diagnosis. The following data were collected at the enrollment visit: patient characteristic and vascular pythiosis-related data (anatomical lesions, disease duration, type of surgery, antifungal treatment, and iron chelating therapy). During the follow-up course, signs and symptoms of possible residual disease including fever, pain, skin rash, mass at surgical sites, arterial insufficiency syndrome, inflammation at surgical site were recorded (Table 1).

**Table 1.** Characteristics of vascular pythiosis patients.

|  | Survived (45) | Deceased (5) | *P*-value |
| --- | --- | --- | --- |
| **Patient related parameters** |  |  |  |
| Age (yr) | 33.6 ± 10.7 | 49.2 ± 17.2 | 0.006 |
| Male sex | 25 (56%) | 4 (80%) | 0.29 |
| Occupation |  |  | 0.39 |
| - Agriculture related | 39 (86.7%) | 5 (100%) |  |
| - Non-agriculture related | 6 (13.3%) | - |  |
| History of water exposure within 3 months | 41 (91.1%) | 5 (100%) | 0.49 |
| Underlying disease |  |  | 0.87 |
| - α-thalassemia | 3 (6.7%) | - |  |
| - β-thalassemia | 3 (6.7%) | - |  |
| - β-thalassemia Hemoglobin E disease | 32 (71.1%) | 4 (80%) |  |
| - H-constant spring | 2 (4.4%) | - |  |
| - Hemoglobin H disease | 5 (11.1%) | 1 (20%) |  |
| Serum ferritin (ng/mL) | 1,388.40 ± 653 | 1,676 ± 402.4 | 0.34 |
| Period from onset of disease to first medical attention (mo) | 1.9 ± 0.7 | 4. ± 0.7 | <0.0001 |
| Period from diagnosis to first definitive surgery (mo) | 0.6 ± 0.2 | 1.5 ± 1.2 | <0.0001 |
| **Disease and treatment related parameters** |  |  |  |
| Location of lesions |  |  | 0.05 |
| - Brachial artery | 1 (2.2%) | - |  |
| - Radial artery | 11 (24.4%) | - |  |
| - Ulnar artery | 1 (2.2%) | - |  |
| - Femoral artery | 17 (37.8%) | 4 (80%) |  |
| - Anterior tibial artery | 9 (20%) | - |  |
| - Posterior tibial artery | 3 (6.7%) | - |  |
| - Iliac artery | 3 (6.7%) | - |  |
| - External carotid artery | - | 1 (20%) |  |
| Surgical procedures |  |  | <0.0001 |
| - Amputation | 45 (100%) | 2 (40%) |  |
| - Debridement | - | 3 (60%) |  |
| Antifungal agents |  |  | 0.17 |
| - Itraconazole alone | 5 (11.1%) | 2 (40%) |  |
| - Itraconazole + terbinafine | 34 (75.6%) | 3 (60%) |  |
| - SSKI + terbinafine | 6 (13.3%) | - |  |
| Duration of antifungal treatment (mo) | 5.9 ± 4.6 | 1.4 ± 0.9 | 0.04 |
| Iron chelation drug | 45 (100%) | 5 (100%) | - |
| **Clinical signs/symptoms post treatment initiation** |  |  |  |
| - Fever >38.2°C | - | 3 (60%) | 0.008 |
| - Arterial insufficiency syndrome (claudication, paresthesia, gangrenous ulceration) | - | 2 (40%) | 0.008 |
| - Mass at surgical sites (arterial aneurysm) | - | 1 (20%) | 0.1 |
| - New skin lesions | - | - | - |
| - Inflammation/infection at surgical sites | - | 3 (60%) | 0.008 |

Patients were followed up for one year after the primary dose of PIAI and their follow-up visits were synchronized with their PIAI schedule as described below. If patients were transferred back to their home provinces, PIAI was sent to their local hospitals and the case record forms were filled out by their primary care providers. At each follow-up visit, patients’ blood samples were collected by using acid citrate dextrose (ACD) vacuum tubes prior to PIAI administration and the samples were delivered back to KCMH within 24 hours. Patients’ sera were prepared from their blood samples by centrifugation at 1,500 g for 15 minutes and kept at − 80°C. Age and sex matched negative control sera obtained from thalassemia donors without active bacterial or fungal infections were used as control. All control sera were used as negative control for BG and *Pi*-Ab testing. To prove the specificity of *Pi*-Ab assay based on the KCMHELISA, a total of 30 non-pythiosis patients’ serum samples with nocardiosis or other fungal infections (candidiasis, cryptococcosis, aspergillosis, talaromycosis, histoplasmosis), proven by nucleic acid sequence analysis, were tested in parallel as negative control.

### *P*. *insidiosum* immunotherapy preparation and schedule

PIAI was prepared according to the method described by Mendoza L. *et al* (12). Five-day-old *P*. *insidiosum* isolates were cultured in Sabouraud dextrose broth at 37°C and inactivated by 0.02% (wt/vol) Thimerosal for 30 min. The antigens were precipitated from disrupted mycelial masses and culture supernatant. For long-term storage, obtained antigens were lyophilized and kept at –80°C until used. 1 mL of 2 mg/mL PIAI was administered via subcutaneous route according to PIAI schedule: The first dose (time 0) was administered as soon as the definitive diagnosis was established; subsequently, 6 booster doses were administered at 0.5, 1, 1.5, 3, 6 and 12 months.

### Beta-D-glucan assay

Serum BG was quantitated, in duplicate, by using the Fungitell^®^ assay (associates of Cape Cod, Inc., Falmouth, MA, USA) according to the manufacturer’s instructions. A pooled serum of 40 thalassemia, otherwise healthy, volunteers was run in parallel as a negative control. The range of the Fungitell^®^ assay was 7.812 pg/mL–523.438 pg/mL. Samples with BG levels out of the indicated range were reported as <7.812 pg/mL or >523.438 pg/mL and processed in the statistical analyses as 7.812 pg/mL or 523.438 pg/mL, respectively. Additional dilutions were not performed. Serum BG level <60 pg/mL, 60-79 pg/mL, ≥80 pg/mL were interpreted as negative, indeterminate, and positive, respectively (13, 14).

### *P*. *insidiosum* specific antibody assay

Serum *Pi*-Ab levels were monitored by the KCMH-ELISA in quadruplicate. No cross reactions with other bacterial or fungal pathogens by this KCMH-ELISA were observed. Microtiter plates (Polysorp^®^, Nunc, Rochester, NY) were coated with 0.1 mL of 2 mg/mL PIA in bicarbonate buffer, overnight at 4°C, then washed with 1%(vol/vol) Tween20 in phosphate-buffered saline (1% PBS-T; PBS: 137 mM NaCl, 2.7 mM KCl,10 mM Na_2_HPO_4_ and 2 mM KH_2_PO_4_) and blocked with 2% Skim milk in PBS-T, 1 h at 37°C. After washing, total of 100 μL of serum in 0.5% skim milk was added to each well, 1 h at 37°C. Horseradish peroxidase-conjugated rabbit anti-human immunoglobulin G (Dako, Denmark) in PBS-T and O-phenylenediamine dihydrochloride in citric acid with 30% H2O2 were used as secondary antibody and substrate, respectively. Reactions were quenched by 200 μL of 1 N H_2_SO_4_. Optical density (OD) was measured at _490_ nanometers. The pooled serum of thalassemia donors was analyzed in parallel as negative control (15).

#### Optimal serum dilution for *Pi*-Ab assay

The optimal serum titer was defined as the titer that presented the best discrimination of OD_490_ value among pythiosis sera at each PIA injection time-point. Pooled serum samples of five vascular cases, randomly selected from each region in Thailand, were tested. Two-fold serial dilution (1:50 to 1:25,600) of those pooled sera were tested by ELISA in quadruplicate. The correlation between OD_490_ value and serum dilution was plotted using GraphPad Prism 5 program (GraphPad Software, Inc., La Jolla, USA).

#### *Pi*-Ab assay result analysis

To standardize the ELISA result for each testing batch, the OD_490_ values were normalized to “ELISA value (EV)” according to the following formula: EV (OD_sample_–OD_background_)/(OD_control_-OD_background_).

### Statistical analyses

Statistical analyses were conducted by SAS version 9.4 (SAS Institute, Cary, NC, USA). The t-test and the Wilcoxon rank sum test were used to compare continuous covariates between groups of patients who survived and died during the follow-up period. The chi-square test and the Fisher’s exact test were used to compare categorical and binary covariates between the 2 groups.

The linear mixed effect models were used to compare the differences in BG levels and *Pi*-Ab levels among groups of patients who survived and died during the study period. The linear mixed effect model is a regression technique for multiple observations for each individual to allow subset of the regression parameters to vary randomly from one individual to another, thereby accounting for sources of natural heterogeneity of the patients. In addition, this regression method can overcome unbalanced data as some patients passed away at 1.5 months; therefore, no additional BG and *Pi*-Ab levels from those patients was available for analysis beyond the 1.5-month mark. We ran the regression models to compare BG and *Pi*-Ab levels between the 2 groups for up to 3 months of the follow-up period. In this analysis, we used unstructured covariance matrix in the regression models.

### *In vitro* susceptibility testing

*In vitro* susceptibility tests were performed with zoospores of *P*. *insidiosum*, isolated from patients with positive cultures, according to the CLSI M38-A2 protocol (16). All *P*. *insidiosum* isolates were confirmed by the PCR methods. *Candida parapsilosis* (ATCC22019) and *Aspergillus flavus* (ATCC204304) were used as controls. All isolates were tested against 7 antifungal agents: amphotericin B, voriconazole (VRC), itraconazole (ITC), fluconazole, anidulafungin, caspofungin, and terbinafine (TRB). The MICs of these antifungal agents ranged from 0.125 to 64 mg/L. Synergistic effects of VRC/TRB and ITC/TRB were tested by the checkerboard technique. The MICs of the individual agents and combination drugs were interpreted in the unit of mg/L and the fractional inhibitory concentration index (FICI) according to the following formula: FICI = (MIC_TRB in combination_/MIC_TRB alone_)+(MIC_VRC or ITC in combination_/MIC_VRC or ITC alone_), respectively. A FICI ≤0.5, >0.5–4.0, >4.0 indicated synergistic, indifferent and antagonistic effect, in order. 0.05% benomyl was used as a positive control.

## Results

### Patient characteristic

Fifty patients met the diagnostic criteria for vascular pythisosis and were recruited in this study. Twenty-two patients had positive culture, 28 patients had positive PCR-ITS/*COX2* results, and 50 patients had positive serum *Pi*-Ab. During the study period, 45 patients survived and 5 patients died. Baseline patient characteristics, treatment modalities, and post treatment clinical information were summarized in Table 1. Definitive surgeries, defined by achievement of negative surgical margins, were achieved in all patients in both groups except one patient with a carotid lesion in the deceased group, where the definitive surgery was not possible. Three of non-survivors underwent debridement to save their limbs. Duration of antifungal therapy in the deceased group was significantly shorter than in the survival group (mean duration 1.4±0.9 vs 5.9±4.6 months; p=0.04); however, this was because patients in the deceased group did not live long enough to complete the therapy.

Among the five patients who died, the patient with carotid disease was determined not to undergo further evaluation. Only one out of four in the deceased group, who initially had femoral disease, was further evaluated based on the clinical symptoms. Three of the patients were initially diagnosed with surgical wound infection, however, they were ultimately found to have residual pythiosis and died 3 months after enrollment. None of patients were on hemodialysis or received intravenous amoxicillin/clavulanate, albumin, intravenous immunoglobulin, potential sources of contaminating BG, during the study period.

### Serum BG levels among survivors and deceased patients

At the time of diagnosis, the means of BG level in survival group and deceased group were not statistically different (489.5±39.6 pg/ml vs 514.3±14.0 pg/ml respectively; p=0.17). After the first dose of PIAI, patients in the survival group had significantly lower means of serum BG than the deceased group (Table 2). Based on the recommended cut-off value, >80 pg/mL, for serum BG, all patients in this cohort had positive test at the time of diagnosis, whereas, all results from thalassemia donors (control group) were negative (Figure 1).

**Table 2.** Mean (SD) serum β-D-glucan level (pg/mL) comparing deceased and survived groups

| Time (mo) | Deceased; n = 5 | Survived; n = 45 | *P-values* |
| --- | --- | --- | --- |
| 0 | 514.35 (14.0) | 489.57 (39.6) | 0.17 |
| 0.5 | 518.70 (4.7) | 387.68 (55.4) | < 0.001 |
| 1.0 | 511.49 (8.5) | 298.80 (46.9) | < 0.001 |
| 1.5 | 493.76 (30.6) | 154.56 (33.9) | < 0.001 |
| 3.0 | 471.67 (24.7)* | 80.30 (22.5) | < 0.001 |

\*only 3 patients

**Figure 1.**
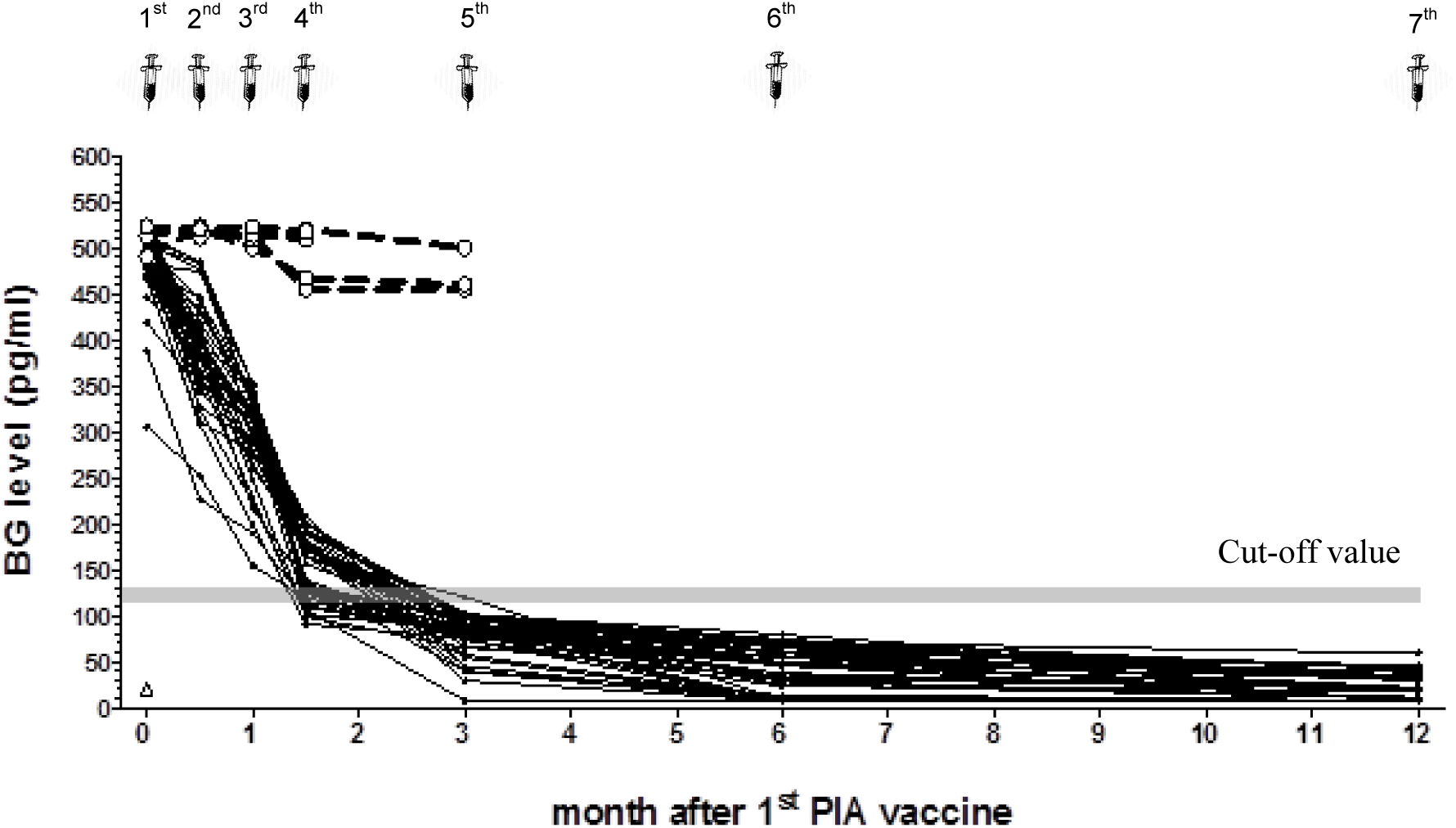
The individual values of serum beta-D-glucan (BG) levels of vascular pythiosis patients who survived (n=45 (•,—)) and died (n=5 (O,‐‐‐)) during 1 year follow-up along with *Pythium insidiosum* antigen immunotherapy (PIAI) treatment 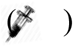 and control (pooled sera of thalassemia group; n=40 (Δ)). Serum BG determined by Fungitell^®^ assay (Associates of Cape Cod, Inc., East Falmouth, MA, USA).

At each follow-up visit during the first 3 months, the means of serum BG levels significantly deceased among the survived group and became negative after 3 months (Figure 1). However, the means of serum BG levels in deceased group did not significantly change during the follow-up period (Table 2). In fact, the mean of BG levels at 3 months remained highly positive until patients died.

### *P*. *insidiosum* specific antibody among survival and deceased patients

By *in*-*house* ELISA assay, the serum dilution of 1:800 presented the most significant difference between upper and lower limitation of OD_490_ value at each PIAI time-points (Figure 2). Therefore, 1:800 dilution was used for the tested sera in this project.

**Figure 2.**
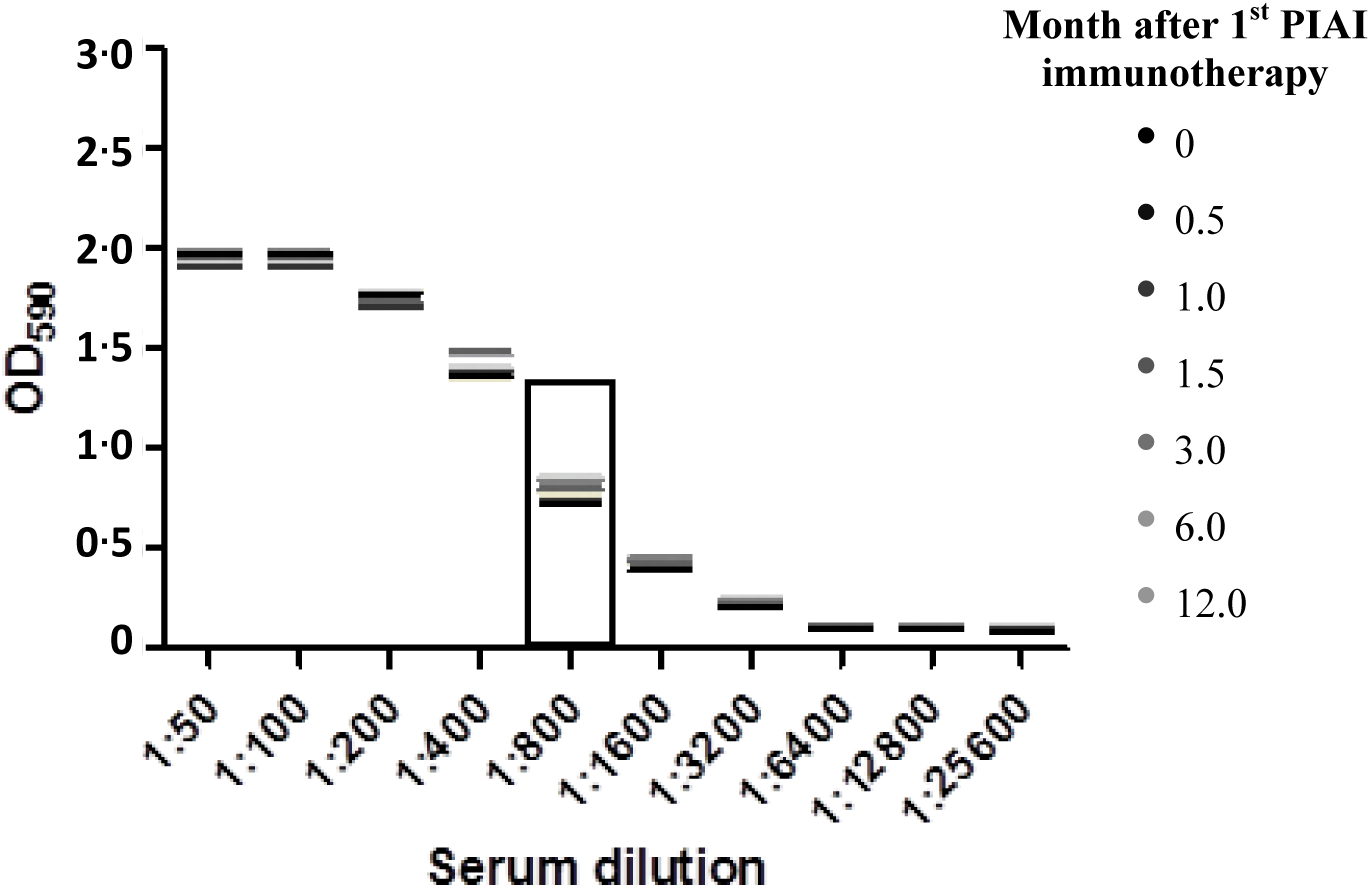
Comparison of OD_490_ value obtained from 7 sera collected before each PIAI time points by using in-house enzyme-linked immunosorbent assay (ELISA) assay. Each serum sample was pooled from randomly selected of five patients. All sera samples were performed by the 2-fold serial dilution from 1:50 to 1:25,600 (X-axis). Each line demonstrated the mean of OD_490_ value (Y-axis).

Based on the results from the linear mixed effect model (Table 3), the mean EV value of *Pi*-Ab in the survival group was significantly higher than in the deceased group, at diagnosis (8.21±0.7 and 2.43±0.2 respectively; p<0.001). At each follow-up visit, there were no significant changes in EV value of *Pi*-Ab levels among survivors. Patients in the survived group maintained their EV value of *Pi*-Ab above 8 throughout the study period (Table 3, Figure 3). The *Pi*-Ab levels among patients in the deceased group significantly increased by 0.28 per half month (p=0.02). However, the average EV value of *Pi*-Ab remained below 4 during their follow-up period.

**Table 3.** ELISA value of *Pythium insidiosum* antibody in survival and deceased groups

|  | Deceased<br>n = 5 | *P-values* | Survived<br>n = 45 | *P-values* |
| --- | --- | --- | --- | --- |
| Means (SD) of *Pythium* antibody level at diagnosis | 2.43 (0.2) | - | 8.21 (0.7) | <0.001 |
| Rate of change in *Pythium* antibody level per 0.5 mo | 0.28 | 0.02 | 0.29 | 0.95 |

**Figure 3.**
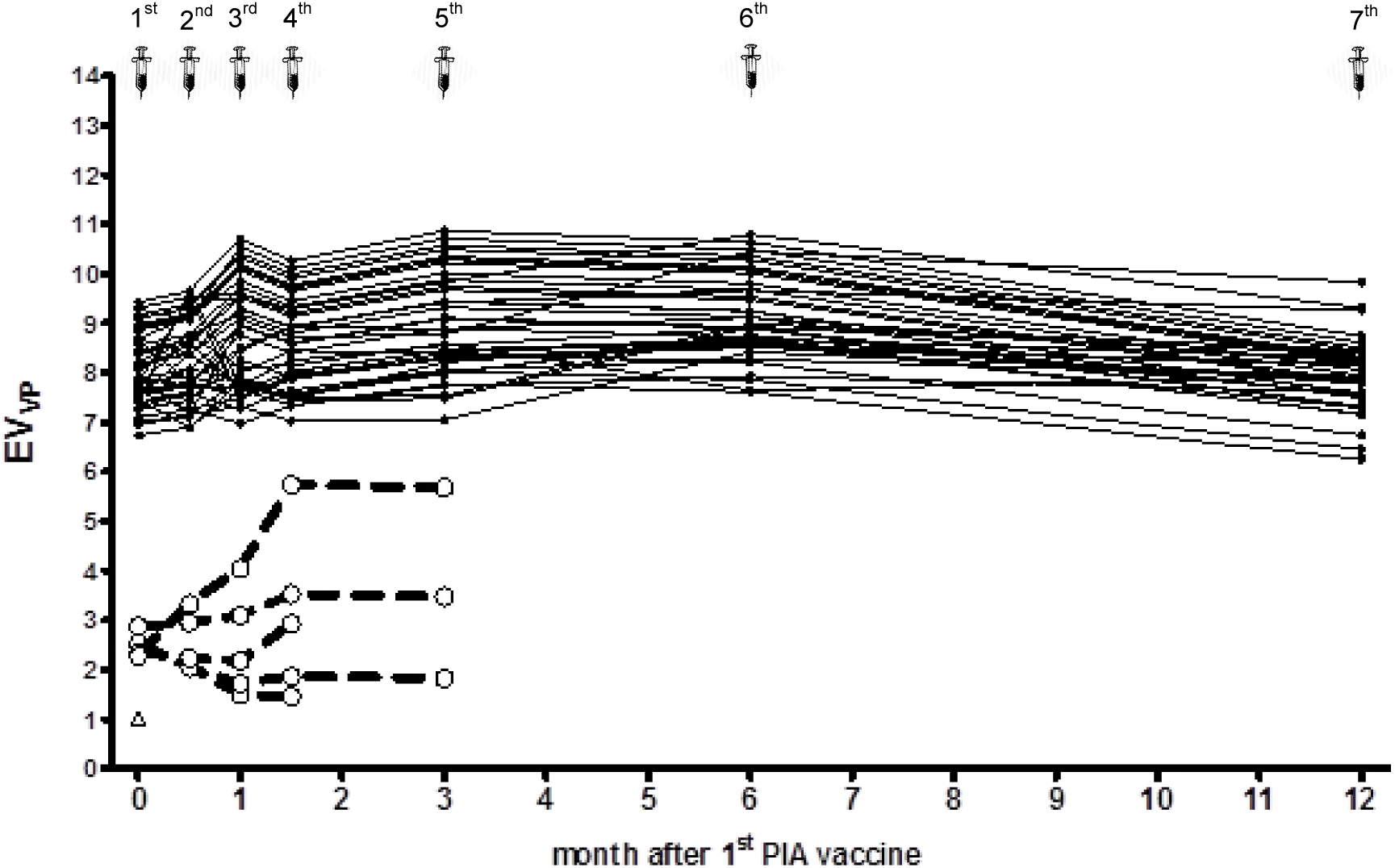
The individual value of antibody response of *Pythium insidiosum* antigen immunotherapy (PIAI)-treated pythiosis patients who survived; n=45 (•,—) and died; n=5 (O,‐‐‐) during 1 year follow-up along with PIAI treatment and control (pooled sera of thalassemia group; n=40 (Δ)) measured by *in*-*house* enzyme-linked immunosorbent assay (ELISA) assay.

### *In vitro* susceptibility results

The highest MICs were observed in AMB, ranging from 4 to 8 mg/L, and the lowest MICs were observed in ITC, ranging from 1 to 4 mg/L. No synergistic effect was found in neither the combination of VRC/TRB nor ITC/TRB with FICI ranging from 1.5 to 2.0 (Table 4).

**Table 4.** MICs of individual and combined agents against *Pythium insidiosum* isolates (n=22)

| Isolate | Patient status | Individual agents (mg/L)<sup>a</sup> |  |  |  |  |  |  | Combined agents (mg/L)<sup>a</sup> |  |  | Combined agents (mg/L)<sup>a</sup> |  |  |
| --- | --- | --- | --- | --- | --- | --- | --- | --- | --- | --- | --- | --- | --- | --- |
|  |  | AMB | VRC | ITC | FLC | ANF | CAS | TRB | VRC | TRB | FICI<sup>b</sup><br>(interpretation) | ITC | TRB | FICI<sup>b</sup><br>(interpretation) |
| 1 | survived | 4 | 2 | 1 | 1 | 2 | 2 | 2 | 1 | 2 | 1.5 (I) | 1 | 2 | 2.0 (I) |
| 2 | survived | 4 | 4 | 2 | 2 | 4 | 4 | 4 | 2 | 2 | 1.0 (I) | 2 | 2 | 1.5 (I) |
| 3 | survived | 8 | 2 | 2 | 4 | 8 | 2 | 2 | 2 | 2 | 2.0 (I) | 2 | 2 | 2.0 (I) |
| 4 | survived | 4 | 4 | 1 | 2 | 8 | 4 | 2 | 2 | 2 | 1.5 (I) | 0.5 | 2 | 1.5 (I) |
| 5 | survived | 8 | 2 | 4 | 8 | 8 | 4 | 4 | 2 | 4 | 2.0 (I) | 2 | 4 | 1.5 (I) |
| 6 | survived | 4 | 2 | 2 | 4 | 8 | 4 | 2 | 1 | 2 | 1.5 (I) | 1 | 2 | 1.5 (I) |
| 7 | survived | 4 | 4 | 4 | 4 | 4 | 2 | 4 | 2 | 4 | 1.5 (I) | 1 | 2 | 1.5 (I) |
| 8 | survived | 4 | 2 | 2 | 2 | 4 | 2 | 2 | 1 | 1 | 1.0 (I) | 2 | 2 | 2.0 (I) |
| 9 | survived | 4 | 4 | 2 | 2 | 2 | 4 | 2 | 2 | 2 | 1.5 (I) | 0.5 | 1 | 1.0 (I) |
| 10 | survived | 8 | 8 | 2 | 2 | 4 | 8 | 4 | 4 | 4 | 1.5 (I) | 2 | 4 | 2.0 (I) |
| 11 | survived | 4 | 2 | 1 | 4 | 4 | 2 | 2 | 1 | 2 | 1.5 (I) | 1 | 2 | 2.0 (I) |
| 12 | survived | 8 | 4 | 2 | 4 | 4 | 2 | 2 | 2 | 2 | 1.5 (I) | 1 | 2 | 2.0 (I) |
| 13 | survived | 4 | 2 | 2 | 2 | 4 | 2 | 4 | 2 | 2 | 1.5 (I) | 1 | 2 | 1.5 (I) |
| 14 | survived | 4 | 4 | 2 | 2 | 4 | 4 | 4 | 2 | 2 | 1.0 (I) | 2 | 2 | 1.5 (I) |
| 15 | survived | 4 | 2 | 4 | 2 | 8 | 4 | 2 | 2 | 2 | 2.0 (I) | 2 | 2 | 1.5 (I) |
| 16 | survived | 4 | 4 | 4 | 2 | 4 | 4 | 4 | 2 | 4 | 1.5 (I) | 4 | 4 | 2.0 (I) |
| 17 | survived | 4 | 4 | 2 | 4 | 4 | 2 | 2 | 2 | 2 | 1.5 (I) | 2 | 2 | 2.0 (I) |
| 18 | survived | 8 | 4 | 2 | 4 | 2 | 4 | 4 | 1 | 4 | 1.25 (I) | 1 | 4 | 1.5 (I) |
| 19 | deceased | 4 | 1 | 4 | 2 | 4 | 4 | 2 | 0.5 | 2 | 1.5 (I) | 2 | 2 | 1.5 (I) |
| 20 | deceased | 4 | 2 | 4 | 4 | 8 | 2 | 4 | 2 | 4 | 2.0 (I) | 4 | 4 | 2.0 (I) |
| 21 | deceased | 8 | 2 | 2 | 2 | 4 | 2 | 2 | 1 | 2 | 1.5 (I) | 2 | 2 | 2.0 (I) |
| 22 | deceased | 4 | 4 | 1 | 4 | 4 | 2 | 2 | 2 | 2 | 1.5 (I) | 1 | 2 | 2.0 (I) |

<sup>a</sup> Concentrations of tested antifungal agents ranged from 0.125 to 64 mg/L.
<sup>b</sup> I, indifferent; FICI, fractional inhibitory concentration index
AMB, amphotericin B; VRC, voriconazole; ITC, itraconazole; FLC, fluconazole; ANF, anidulafungin; CAS, caspofungin; TRB, terbinafine

## Discussion

We describe the first prospective cohort of vascular pythiosis patients who were followed up for one year after diagnosis. In our study, definitive surgery was performed in 49/50 (98%) patients. This is likely the reason why the mortality rate in our study decreased to 10% from 36.4%-44.4% in the previous studies (2, 3, 17). As expected, patients who survived had a significant shorter mean duration from the onset of disease to their first medical encounters as well as a shorter mean waiting time for their definitive surgeries. This emphasizes the importance of early disease detection and prompt surgical intervention to improve the survival chance. During the first 3 months of treatment course, only non-survivors developed fever, inflammation of the surgical sites, mass at the surgical sites, or arterial insufficiency syndrome. Unfortunately, these symptoms are not specific and only one patient was further evaluated, based on these clinical symptoms in a timely manner.

We found all patients had high positive serum BG at the diagnosis. Patients who survived had a significant decrease in BG levels at every visit. Conversely, patients who died had persistently high BG levels throughout the study period. These findings suggest that BG trend may be more important than the level itself. Vascular pythiosis patients with persistently elevated BG should be further evaluated for any residual disease despite documented negative surgical margins from the pathology reports. The use of BG trends has been reported in previous studies of patients with candidemia (18) and *Pneumocystis jirovecii* pneumonia (19). As BG is not a *P*. *insidiosum*-specific biomarker, vascular pythiosis patients with persistent BG elevation should be evaluated for other possible causes of positive BG tests including *Pseudomonas* infection, blood/albumin infusion, cellulose membranes in hemodialysis, and in particular intravenous amoxicillin-clavulanate, which is available in Thailand (20-22).

Similar to BG levels, *Pi*-Ab was positive in all patients in this study at diagnosis based on the cut-off level of EV>1.5. Interestingly, all patients in the survival group were able to maintain their EV values of *Pi*-Ab>8 during the study period, whereas all patients in the deceased group had EV values of *Pi*-Ab<4 throughout the study, despite the PIAI administration per protocol. We suspect that the persistent *Pi*-Ab in surviving patients, without evidence of ongoing infections for a year, is likely due to PIAI administration and this phenomenon has been reported in the literature (6). These findings suggest the EV value of *Pi*-Ab>8 may be a good prognostic indicator that implies good host immune response to PIAI against pythiosis.

Importantly, our results have provided the opportunity to evaluate our treatment protocols. Our data suggest that aggressive surgery is still crucial as the mortality rate in our study remained at 10% despite obtaining negative surgical margins in 98% of the cases. Likewise, we have learned that achievement of negative surgical margins does not necessarily mean patients do not have residual infection. Complaints of any symptoms at the surgical site or proximal vascular lesions should be treated as ongoing vascular pythiosis until proven otherwise.

There are several limitations in this study. Pythiosis is a dangerous infection requiring aggressive therapy and not all patients received exactly the same treatment. Accordingly, it was not possible to control all factors that could affect BG and *Pi*-Ab levels. In addition, as we are studying a relatively uncommon disease, a small sample size is unavoidable. Despite its small sample size, this is still the largest prospective study in vascular pythiosis to preliminarily evaluate BG and *Pi*-Ab levels as potential markers for disease monitoring. This study also adds *in vitro* susceptibility results from human isolates as well as describes clinical outcomes at the current age into the literature. Multicenter prospective studies are still required to determine how to incorporate BG and *Pi*-Ab levels into clinical decision making. In addition, other non-specific inflammatory markers, i.e. c-reactive protein, erythrocyte sedimentation rate, etc., should be investigated as potential lower cost markers.

In summary, we believe that the BG and *Pi*-Ab are the potential markers in the management of vascular pythiosis and should be further investigated to determine whether such monitoring has an impact on clinical outcomes. Persistently elevated BG after a definitive surgery should prompt further evaluation for possible residual disease. The high EV level of *Pi*-Ab may indicate a good host immune response to *P*. *insidiosum* and PIAI.

## Acknowledgements

This work was supported by the 90^th^ and 100^th^ Anniversary of Chulalongkorn University Scholarship; the National Research Council of Thailand (No. 406-699752-4 and FY 2017-Thesis Grant for Doctoral Degree Student), the National Research Council of Thailand and Health Systems Research Institute (No. GRB_BSS_81_59_30_22). NW, RR, AL, AC received the personal fees from The National Research Council of Thailand and Health Systems Research Institute (No. GRB_BSS_81_59_30_22). The funders had no role in study design, data collection and interpretation, or the decision to submit the work for publication. MF is Director, Clinical Development at Associates of Cape Cod, Inc. (Fungitell assay manufacturer). No other conflict of interest to declare.

The authors would like to thank all vascular surgeons, infectious disease specialists and research coordinators at KCMH. We would like to give special thanks to Ms. Supaporn Amornsirivat, Hematology Unit, Faculty of Medicine, Chulalongkorn University for coordinating and supporting this project.

This study was partially presented at the 8^th^ Trends in Medical Mycology (TIMM) at Belgrade, Serbia. (Abstract No. P114).

